# Comparative transcriptome in rhesus macaques and crab-eating macaques

**DOI:** 10.1101/2023.08.17.553631

**Authors:** Yuxiang Mao, Yamei Li, Zikun Yang, Ning Xu, Shilong Zhang, Xuankai Wang, Xiangyu Yang, Qiang Sun, Yafei Mao

## Abstract

Understanding the variations in gene expression between species is pivotal for deciphering the evolutionary diversity in phenotypes. Rhesus macaques and crab-eating macaques serve as crucial nonhuman primate biomedical models with different phenotypes, but the large-scale of comparative transcriptome research between these two species has yet to be fully elucidated. Here, we conduct systematic comparisons utilizing newly sequenced RNA-seq data from 84 samples encompassing 14 common tissues. Our findings reveal that a small fraction of genes (∼3.7%) show differential expression between the two macaque species, while ∼36.5% of genes show tissue-specific expression in both macaques. We also compare gene expression between macaques and humans and ∼22.6 % of the orthologous genes show differential expression in at least 2 tissues. Moreover, ∼19.41% of genes overlapped with macaque-specific structural variants are more likely to show differential expression between humans and macaques. Of these, *FAM220A* shows elevated gene expression in humans compared to macaques because of lineage-specific duplication. In summary, our study presents a large-scale analysis of the transcriptomes within macaque species and between macaques and humans. These insights into gene expression variations will enhance the biomedical utility of macaque models and contributing to the broader realm of primate genomic studies.

## Introduction

Cross-species transcriptome comparisons play a crucial role in advancing our understanding of gene expression patterns within and between species tissues[1, 2], as well as shedding light on the relationship between genetic variation and differences in gene expression[3]. When applied to primates, such comparisons offer valuable insights into primate evolution, human diseases, and the heterogeneity of primate genomes[4–6].

Among the primate species, the *Macaca* genus of Old World Monkeys holds exceptional significance, owing to its broad geographic distribution and remarkable adaptability as nonhuman primates (NHPs)[7, 8]. Notably, rhesus macaque (*Macaca mulatta*, Mmu) and crab-eating macaque (*Macaca fascicularis*, Mfa) stand out as two extensively studied biomedical models, benefiting from their widespread used in research and the huge efforts of macaque genomics in recent years[9]. However, previous comparative transcriptome in the two macaques have been limited to a few tissues[10, 11], thereby limiting our comprehensive understanding of gene expression patterns and differences within the two vital macaque species.

Macaques are genetically closer to humans, with a split occurring ∼25 million years ago, compared to rodents, which diverged around 70 million years ago from humans[12]. Consequently, macaques exhibit physiological, neurobiological, and disease susceptibility similarities to humans. Despite this, the specific gene expression differences between macaques and humans have remained to be elucidated. To address this gap, we sequence transcriptomes of 84 samples across 14 tissues from both Mfa and Mmu. This comprehensive sequencing effort aims to achieve the following objectives: (1) establish global gene expression patterns and tissue-specific expressed genes in the two species, (2) identify differentially expressed genes (DEGs) between the two macaque species, and (3) explore the DEGs between humans and macaques, and the correlation of DEGs and structural variation.

## Result

### Global Gene Expression Patterns in Two Macaque Species

We generated ∼570 GB of transcriptomic data from 84 tissue samples collected from 8 Mfa and 8 Mmu (Table S1, Table S2). After data filtering, 99.55% of clean reads were retained and each sample showed a proper duplication rate (mean: 32.58%), except for the heart tissues in rhesus macaques (mean_Mmu=_55.05%), potentially owing to the higher content of mitochondrial RNA in heart tissues (mean= 3.98%) (Table S3). All sequencing reads excluding mitochondrial RNA were aligned to the high-quality long-read assembly (rhesus macaque: mmu10) with average alignment rate of ∼88.36% (Table 1)[13].

**Table 1.** Statistics of Macaque Transcriptome Data.

| Average | Sample size |  | Total data(Gbp) |  | Overall Aligned Rate(%) |  | Reads duplicated rate(%) |  |
| --- | --- | --- | --- | --- | --- | --- | --- | --- |
| Tissue/Species | Mfa | Mmu | Mfa | Mmu | Mfa | Mmu | Mfa | Mmu |
| blood | 3 | 3 | 7.01 | 6.57 | 90.59 | 94.63 | 25.65 | 26.62 |
| lung | 3 | 3 | 6.47 | 6.39 | 90.14 | 94.44 | 21.25 | 34.18 |
| liver | 3 | 3 | 6.86 | 7.19 | 88.85 | 92.02 | 29.25 | 52.66 |
| hippocampus | 3 | 3 | 6.41 | 7.45 | 83.38 | 92.69 | 22.25 | 36.92 |
| muscle | 3 | 3 | 6.09 | 7.15 | 84.47 | 95.22 | 30.57 | 42.27 |
| ovary | 2 | 3 | 6.70 | 6.71 | 91.09 | 93.88 | 22.19 | 40.02 |
| spleen | 3 | 3 | 6.16 | 7.14 | 88.64 | 91.98 | 25.36 | 48.65 |
| frontal lobe | 3 | 3 | 6.82 | 7.07 | 81.70 | 92.26 | 26.17 | 37.38 |
| thalamus | 3 | 3 | 6.71 | 7.42 | 82.83 | 93.34 | 24.74 | 37.38 |
| kidney | 3 | 3 | 7.07 | 6.63 | 83.50 | 93.06 | 26.57 | 34.81 |
| visual cortex | 3 | 3 | 6.48 | 6.81 | 79.84 | 90.52 | 26.33 | 41.26 |
| striatum | 3 | 3 | 6.09 | 7.80 | 79.18 | 92.37 | 23.60 | 38.31 |
| cerebellum | 3 | 3 | 6.55 | 7.26 | 85.73 | 94.43 | 23.22 | 32.85 |
| heart | 3 | 4 | 6.47 | 6.58 | 69.80 | 83.40 | 31.09 | 51.68 |

The hierarchical clustering analysis of median gene expression across 14 tissues showed that tissues tended to cluster together, rather than being clustered based on the species. In particular, brain tissues, including neocortical regions (prefrontal and visual cortices), subcortical regions (striatum, hippocampus and thalamus) and cerebellum, showed the most similar expression patterns between the two species and their expression patterns are distinct to other tissues (Fig. 1A). We also observed that low intra-group variability and high reproducibility within the individual groups (Fig. S1). In addition, the number of expressed genes in each tissue was a significantly correlated between the two macaque species (Spearman’s, r^2^ = 0.76, p < 2.2*e*^−16^) (Fig. 1B). Principal component analysis (PCA) provided an additional support, showing that PC1 accounts for the differences between brain tissues and other tissues (Mfa: 34.80% variation, Mmu: 36.64% variation), while PC2 explains the variation among different visceral organs (Mfa: 14.75% variation, Mmu: 13.05% variation) in both species (Fig. 1C).

**Figure 1.**
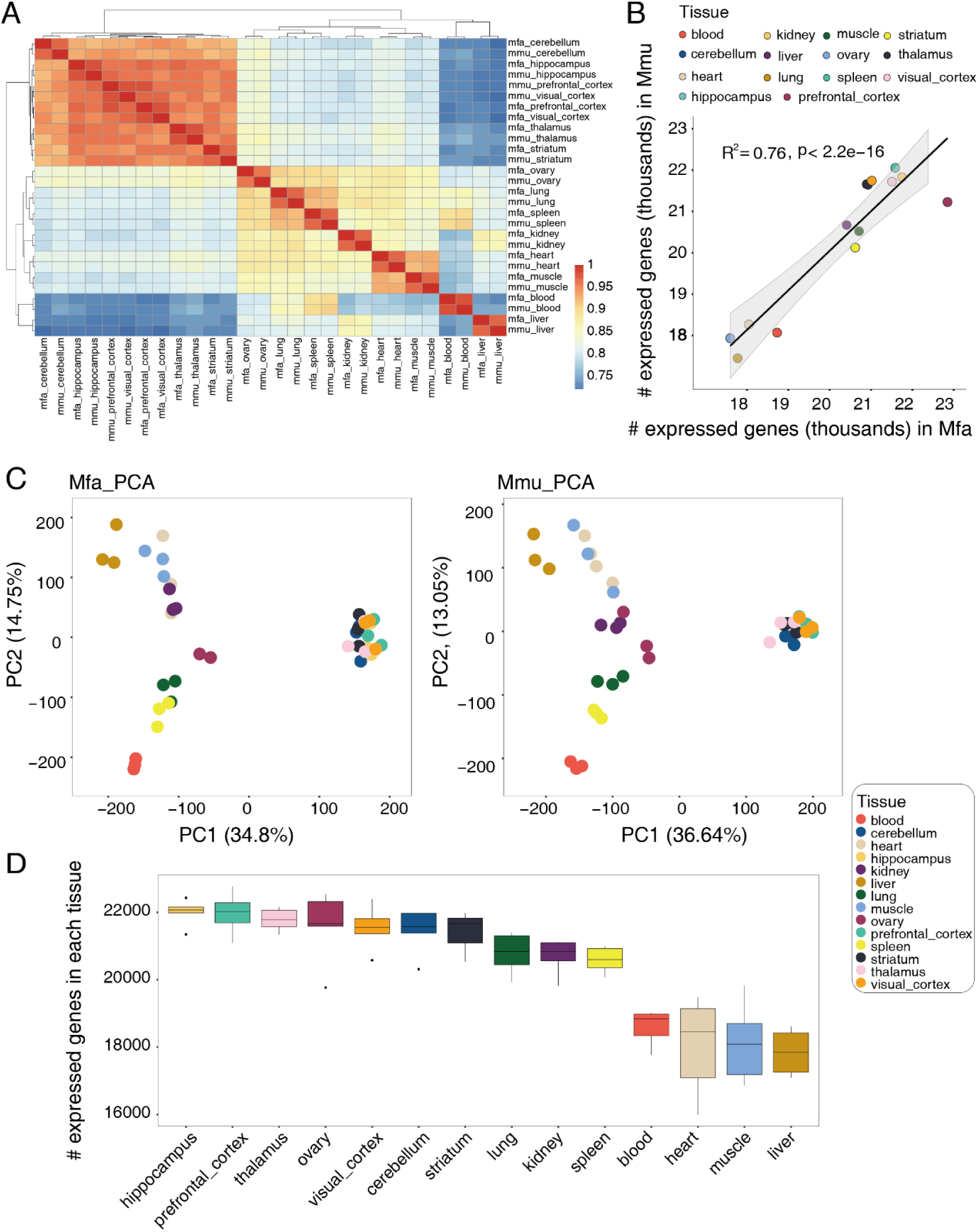
Quality control and global gene expression patterns of macaque tissues. (A) Hierarchical clustering of 14 tissues in crab-eating macaque (Mfa) and rhesus macaque (Mmu) based on Pearson’s correlation of median gene expression value (B) Spearman’s correlation of the number of expressed genes (median TPM > 0.1) between Mfa and Mmu. Gray area represents the 95% confidence interval. Each dot represents a tissue. (C) Principal component analysis of gene expression. Each dot represents a sample, colored by tissue types. The left panel refers to the 41 Mfa, and the right panel refers to the 43 Mmu. (D) The number of expressed genes (TPM > 0.1) identified in different tissues. In each boxplot, the black line represents the median; gray dots represent outliers.

Based on the gene models from a long-read macaque genome assembly (mmu10), we identified 33,634 genes (99.12%) expressed in our dataset. Yet, 349 and 356 genes in Mfa and Mmu, respectively, showed no measurable expression (Transcripts per Kilobase Million, TPM < 0.1) across the 14 examined tissues. Among these, 296 overlapped genes (84.8% vs. 83.1%, Hypergeometric test, p < 2.2×10^-16^) were significantly enriched in male reproductive processes (e.g., spermatogenesis and male gamete generation) (p-adjusted=0.01) as well as defense responses to bacteria (p-adjusted =0.003) and sensory perception (p-adjusted =0.01) (Fig. S2, Table S4). The lack of male samples and skin tissues in this study may contribute to the observation of these expression patterns. As shown in the previous study[14, 15], ovary (#gene mean = 21,586) and brain tissues (#gene mean = 21,714) expressed a greater number of genes, while muscle (#gene mean = 18,086) and liver (#gene mean = 17,845) expressed fewer genes in comparison (Fig. 1D).

### Tissue Specificity of Gene Expression in Two Macaque Species

While Mfa and Mmu showed significantly analogous gene expression patterns, they are distinct species that diverged from each other from ∼2-3 million years ago[16]. Then, we are interested in their gene expression differences. To understand the gene regulation differences between them, we firstly focused on examining the degree of divergence in tissue-specific gene expression. Globally, tissue-specific genes in the same tissues showed significant overlap between the two species. When taking brain tissues individually for comparison, blood tissues showed the highest number of tissue-specific genes across the 14 examined tissues (n_Mfa_ = 2,381, n_Mmu_ = 1,635, n_overlap_=1,309). Yet, the prefrontal cortex showed the least tissue-specific expression (n_Mfa_ = 15, n_Mmu_ = 20, n_overlap_=5) (Fig. 2A). In each tissue, tissue-specific genes were almost identical in the two macaque species (Fig 2B). The number of tissue-specific gene were also significantly correlated in Mfa and Mmu (Spearman’s, r^2^ = 0.92, p < 2.2*e*^−16^) (Fig 2C).

**Figure 2.**
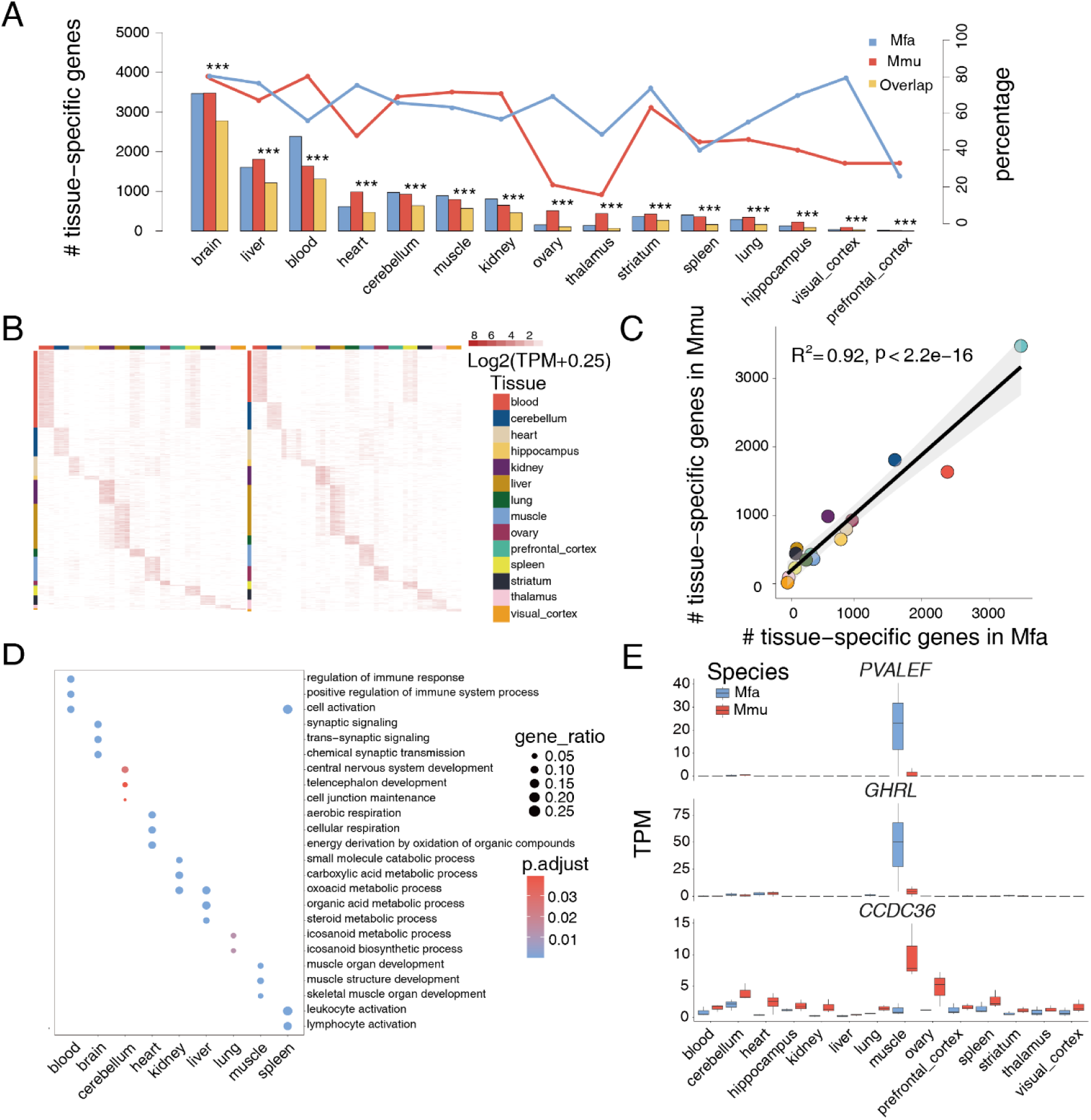
Comparison of tissue specificity of gene expression in macaques. (A) The number of tissue-specific genes (log2(fold-change) > 1.5 and p-adjusted < 0.05) in Mfa (blue) and Mmu (red). The overlapped tissue-specific genes are represented in yellow. “***” represents p-adjusted (Benjamini-Hochberg method corrected p-value) less than 1.0×10^−3^ with hypergeometric test. The lines show the percentage of overlapped tissue-specific genes in each species (blue: Mfa; red: Mmu). (B) Expression profiles of tissue-specific genes in Mfa (left) and Mmu (right). Each row represents a gene and each column represents a sample from the corresponding tissue. (C) Spearman’s correlation of the number of tissue-specific genes across 14 tissues between Mfa and Mmu. Gray area represents the 95% confidence interval. Each dot represents a tissue. (D) Gene ontology (GO) enrichment analysis of overlapped tissue-specific genes. The top 3 most significant functional enrichment terms are shown in each tissue. (E) The expression profiles of *NTRK2*, *SYN1*, and *SNAP25* as brain specific expressed genes.

The relatively similar gene expression patterns were observed in the five different brain tissues and this is a reason why we observed less tissue-specific genes in each brain tissue. Then, we combined five brain tissues for further analysis to identify brain tissue-specific genes. This approach showed a greater number of tissue-specific genes in the brain tissues (n_Mfa_ =3,467, n_Mmu_ = 3,477), with 2,778 genes showing significant overlap between Mfa and Mmu (79.89% and 80.12%). Visceral tissues also showed substantial overlaps, with overlapping rates ranging from 40% to 80%. However, the ovary tissues showed less overlap in comparison (69.23% in Mfa, 21.14% in Mmu) (Fig.2A and S2). One of the reasons for the discrepancy in ovary-specific genes between the two species could be attributed to their evolutionary divergence. Or, the distinct tissue-specific expression in the ovary tissue might stem from differences in the physiological status of the ovaries, such as the estrus cycle phase and ovulation timing. Gene Ontology (GO) enrichment analysis revealed divergent pathway enrichment patterns for ovary-specific genes between the two macaque species (Fig. S3). In Mfa, these genes were enriched in extracellular matrix organization and collagen fibril organization, whereas in Mmu, they were enriched for reproductive pathways like cell division and cell cycle process[17] (Fig. S3).

The tissue-specific genes that overlap between the two species were almost identical as human tissue-specific genes, indicating the conservation across evolutionary lines (Fig. S4). The two different troponin genes: *TNNT2* and *TNNC2*, were specifically expressed in heart and muscle tissues, respectively, in two macaques (Fig. S4). The gene expression patterns were also observed in human expression data. These data support that the tissue-specific genes shared by species (conserved) accurately reflect the known biology of tissues (Fig.2D, Table S5). For example, brain-specific genes of Mfa and Mmu were enriched in chemical synaptic transmission and neuron differentiation (Fig. S5).

However, 3,152 and 2,916 tissue-specific genes were uniquely expressed in Mmu and Mfa, respectively, such as Mfa uniquely expressed gene *PVALEF*, *GHRL* and Mmu uniquely expressed *CCDC36* in muscle tissue (Fig. 2E). Interestingly, Mfa uniquely brain-specific expressed genes were involved in odontogenesis and dentin formation, whereas Mmu uniquely brain-specific expressed were enriched in potassium ion import across the plasma membrane (Table S6).

### Differentially Expressed Genes Between Macaque Species

The DEG analysis between Mfa and Mmu showed the highest number of DEGs in the brain, blood and lung tissues (Fig.3A, Table S7). In contrast, the heart tissues had the lowest number of DEGs, coincident with previous report suggesting slower evolution of heart tissues in mammals[14](Fig. S6). Blood and spleen tissues showed more intra-group variation compared to other tissues (Fig. S7). These variations suggested that differences in gene expression in these tissues may not solely be attributed to species divergence but rather variations in immune status among individuals. Lung tissues also showed significant immune differences (Fig. S8), but the macaques used in this study are devoid of viral infections and no anatomical lesions are observed in lung tissues. Therefore, this led us to suspect the DEGs in lung tissues as potential species-level differences between Mmu and Mfa rather than pathological manifestations. These findings highlight the importance of understanding species-specific immune responses in different macaque species[18, 19].

**Figure 3.**
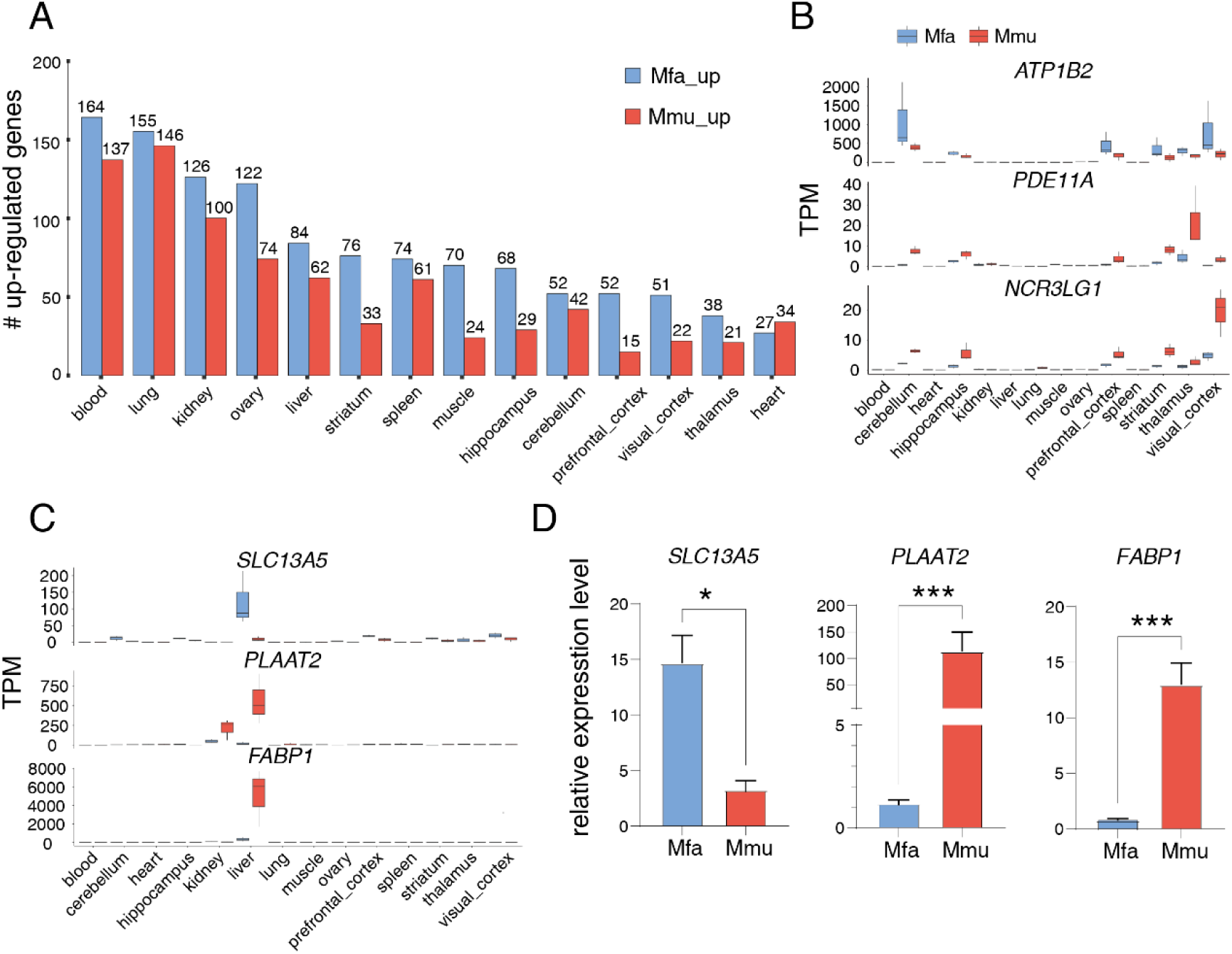
Comparison of gene expression across 14 tissues between two macaques. (A) The number of significantly upregulated genes in Mfa (blue) and Mmu (red) (log_2_FoldChange (log_2_FC)> 1.5 and p-adjust < 0.05). (B) Expression profiles of brain DEGs (*ATP1B2*, *PDE11A*, and *NCR3LG1*) in Mfa(blue) and Mmu (red). (C) Expression profiles of liver DEGs (*SLC13A5*, *PLAAT2*, and *FABP1*) in Mfa (blue) and Mmu (red). (D) Quantitative real-time PCR validation of three liver DEGs (Mfa: n=3; Mmu: n=2). Ctrl: Mfa (*PLAAT2* and *FABP1*), Mmu (*SLC13A5*). Data represents three biological repeats and is shown as mean ± S.E.M. Statistical significance is calculated using wilcoxon test (*p_value < 0.05; ***p_value < 0.001)

For DEGs analysis in brain tissues, we combined all brain tissues firstly and identity 441 Mfa-upregulated and 356 Mmu-upregulated genes, including genes like *ATP1B2*, *PDE11A*, and *NCR3LG1* (Fig. 3B). Notably, some of these genes were related to neurogenic diseases in humans. For instance, *ATP1B2* is involved in maintaining the membrane potential and is down-regulated in schizophrenic patients[20]. The gene expression of *PDE11A* is significantly decreased in brain samples of patients with Alzheimer’s disease, as reported in previous studies[21]. These findings underscore the importance of considering the expression divergence of disease-associated genes in brain tissues when choosing between Mfa and Mmu as a model for specific neurological disorders.

Mfa-upregulated genes in liver and kidney, two metabolism-related tissues, were significantly enriched for amino acid and organic acid transport pathways, while Mmu-upregulated genes were enriched in pathways related to organic hydroxy compound transport (Table S8). These findings indicate that Mfa and Mmu may show some divergences in metabolism. For example, *SLC13A5* involved in citrate metabolic processes was significantly upregulated in the Mfa liver compared to the Mmu liver[22]. In addition, *PLAAT2* and *FABP1,* related to phospholipids and fatty acid metabolic processes[23, 24], respectively, were highly expressed in the Mmu liver but not in the Mfa liver (Fig. 3C). We validated our RNA-sequencing analysis of these three genes with quantitative Reverse Transcription PCR (qRT-PCR), and the qRT-PCR results supported our observations (Fig. 3D). These findings highlight the distinct metabolic differences between Mfa and Mmu, further emphasizing the importance of considering species-specific metabolic variations when using these macaque models in metabolism-related studies.

### Comparative Transcriptomics of Macaques and Humans

To better understand the utility of macaques as biomedical models[25], we are also interested in characterizing the comparative transcriptomics of macaques and humans. Then, we integrated transcriptomes of 84 human samples from the 14 corresponding tissues, downloaded from public databases, with our macaque data for a comparative transcriptomics analysis. Brain, lung, spleen, heart and muscle tissues were clustered by species rather than by their specific tissues (Fig.4A). In contrast, tissues such as ovaries, kidneys, and livers showed clustering based on tissue type rather than species. The correlations observed across the 14 tissues between macaques and humans showed a significant degree of similarity (Wilcoxon test, p=0.91) (Fig. 4B). These findings underscore the potential of macaque models to closely mimic human gene expression patterns across various tissues.

**Figure 4.**
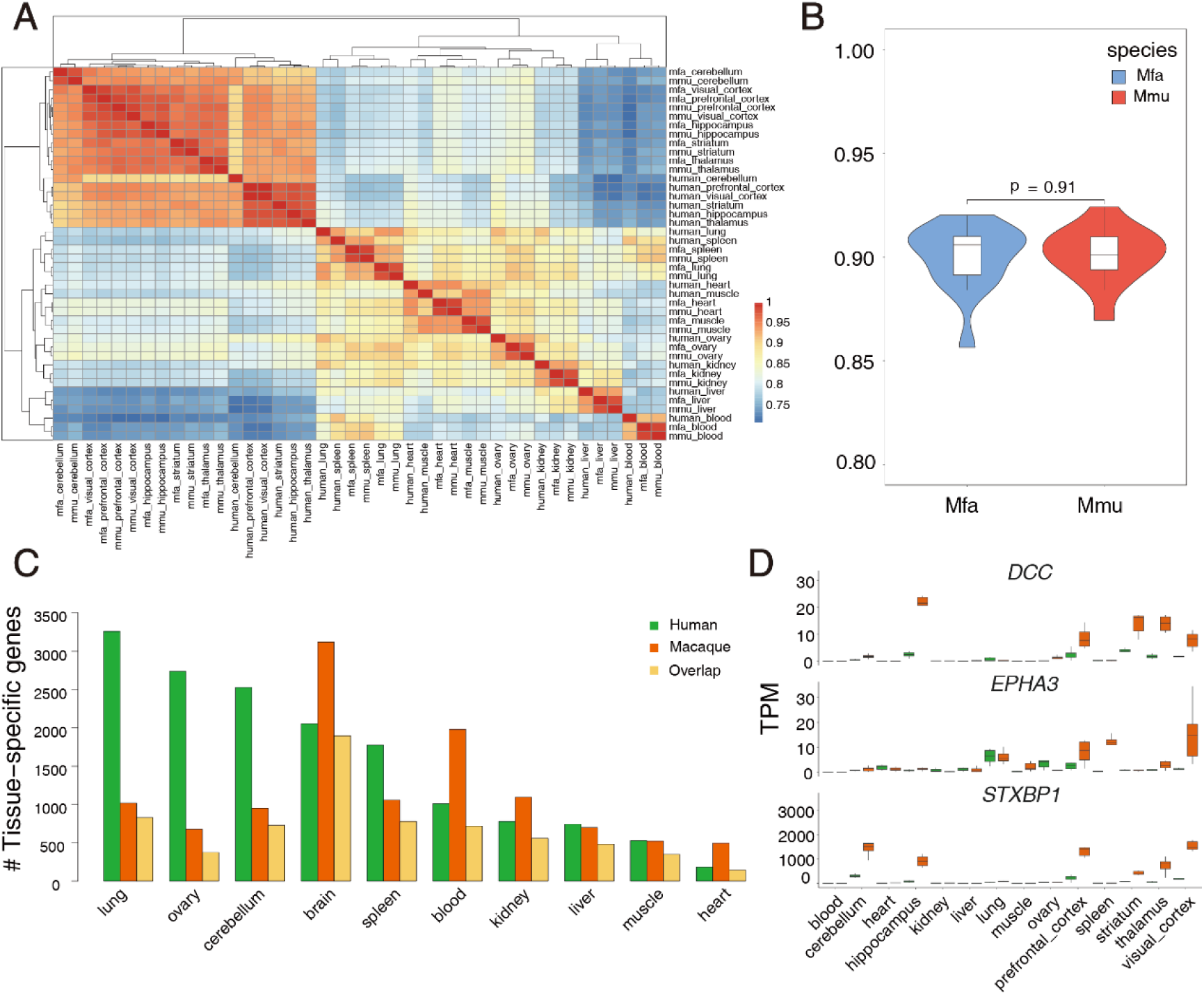
Correlation and tissue specificity in human and macaques. (A) Hierarchical clustering of 14 tissues in humans and macaques based on Pearson’s correlation of median gene expression value. (B) The violin plot shows the correlations of gene expression of tissues between humans and Mfa (blue) and between humans and Mmu (red) (p=0.91, n=14, Wilcoxon test) (C) The number of tissue-specific genes (log_2_FoldChange > 1.5 and p-adjusted < 0.05) and of the overlapped gene in humans (green) and macaques (orange). The overlapped tissue-specific genes are represented in yellow (D) gene expression profile of three human neurological disease-associated genes (*DCC*, *EPHA3*, *STXBP1*) in humans (green) and macaques (orange).

Brain tissues showed the largest number of tissue-specific genes, while heart tissues had the fewest tissue-specific genes both in the two species (Fig.4C), aligning with the patterns observed in the macaque tissue-specific analysis discussed above. The overlapped genes in brain tissues encompassed many critical human neurological disease-associated genes (e.g., *DCC*, *EPHA3* and *STXBP1*) (Fig.4D). *DCC* guides axon growth during neural development and is associated with various neurological diseases[26]. *STXBP1* is crucial for neurotransmitter release, and generation of Mfa carrying *STXBP1* mutation showing focal epilepsy as a primate model of human genetic disorder[27, 28]. These findings further underscore the relevance of macaques as models for studying neurological diseases and advancing our understanding of human health.

Next, we are interested in the gene expression differences between macaques and humans. The analysis of DEGs between macaques and humans showed that brain tissues exhibited the greatest number of differentially expressed genes, while ovary tissues showed the fewest differentially expressed genes (Fig. 5A). From a panel of 15,000 one-to-one orthologous genes between humans and macaques, we identified 7,652 DEGs, of which 3,394 genes shown differential expression in at least two tissues (Table S9). Of these, some were differentially expressed across multiple tissues, including genes like *PSMB2* involved in protein catabolism[29]; *POLR2F* and *KAT14* associated with transcriptional regulation[30, 31]; and *FAM220A* related to positive regulation of protein binding activity (Fig. 5B)[32].

**Figure 5.**
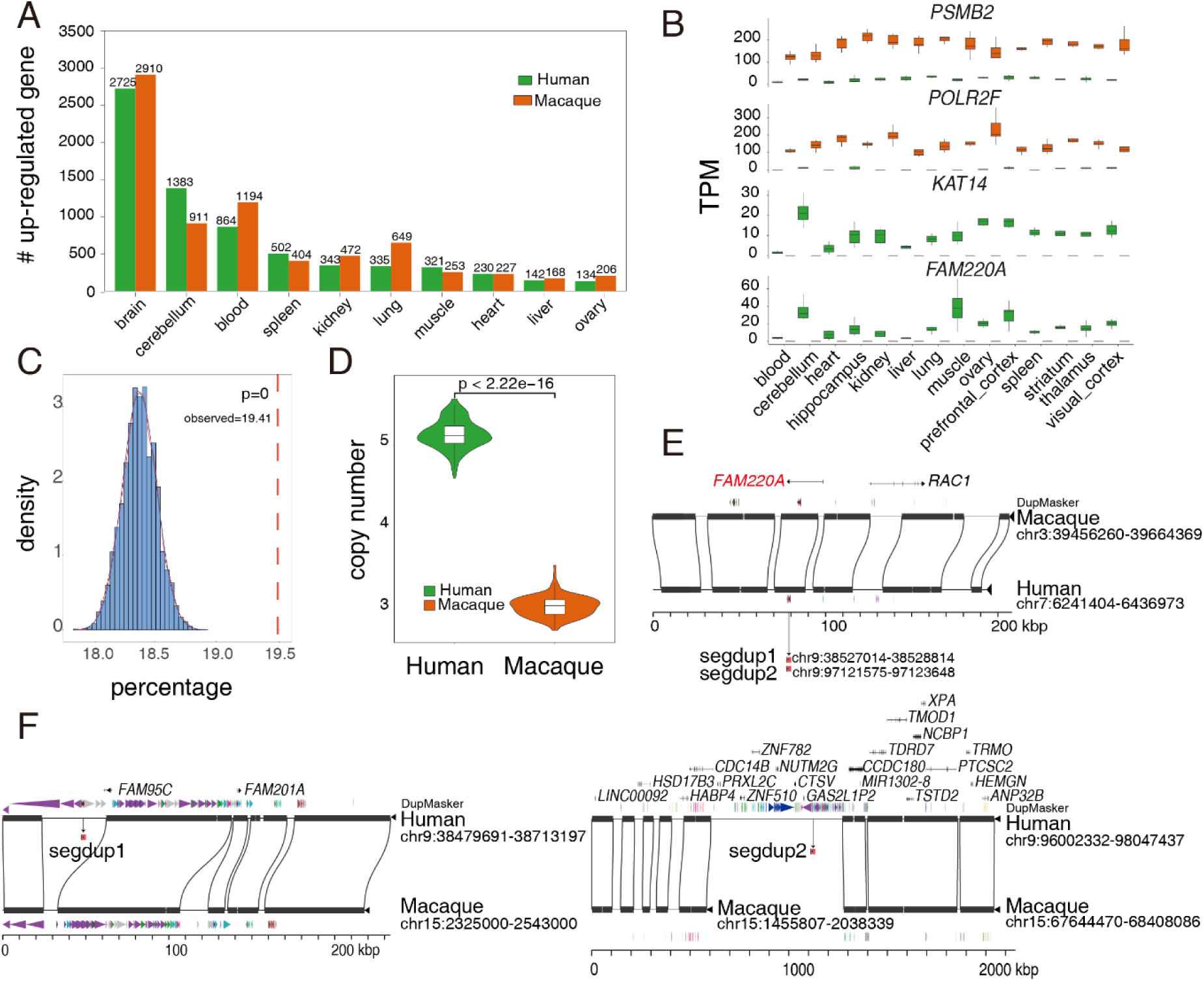
Differential gene expression between humans and macaques. (A) The number of significantly upregulated genes in humans (green) and macaque (orange) (log_2_FoldChange (log_2_FC)> 1.5 and p-adjust < 0.001). (B) Expression profiles of DEGs (*PSMB2*, *POLR2F*, *KAT14* and *FAM220A*) between human (green) and macaque (orange). (C) The distribution of percentage of the intersected DEGs with lineage-specific SVs over the total number of gene intersected with lineage-specific SVs (n=5,000). The red dashed line represents the observed value: the percentage of DEGs intersected with lineage-specific SVs over the number of genes intersected with lineage-specific SVs (empirical p=0). (D) The distribution of copy number of *FAM220A* in humans (green, n=208) and macaques (orange, n=132). (E) Syntenic relationship of *FAM220A* locus between human (GRCh38) and macaque (Mmu10). The gene model, segdup are shown in top and bottom. (F) Syntenic relationship of a human segdup of *FAM220A* locus between human (GRCh38) and macaque (Mmu10) suggest the *FAM220A* duplication in humans.

Then, we investigated the correlation of structural variation and DEGs. We intersected the macaque-specific structural variants (SVs) and the identified DEGs to show that ∼19.41% of genes overlapped macaque-specific SVs are more likely to show differential expression in at least two tissues (n_DEG_=2,601, permutation test, p=0) (Fig.5C). These DEGs were likely enriched in RNA regulation and protein synthesis and metabolism (Fig. S9). In particular, there was a significant difference in the copy number of *FAM220A* between human (mean copy number=5.13) and macaque populations (mean copy number =3.04) (Fig.5D). Further, we found that *FAM220A* has two additional segmental duplications in humans (chr9: 38527014-38528814, chr9: 97121575-97123648), not present in macaques (Fig.5E). This case exemplifies how SVs could impact gene regulation between humans and macaques.

## Discussion

Gaining insight into the overarching patterns of gene expression in two species significantly contributes to our understanding of conservation and heterogeneity between them[14]. In this study, we conduct a comprehensive comparison of transcriptomes across 14 tissues in two macaque species, as well as between humans and macaques. This dataset represents an important initial step in profiling the macaque transcriptome.

Mfa and Mmu are prominent nonhuman primate biomedical models, but they have a few phenotypic differences (e.g., tail length)[33]. Our data illustrates that over ∼96.3% of genes are expressed consistently, and the two macaque species share the majority of tissue-specific genes across the 14 tissues. Notably, our analyses reveal that brain tissues show the highest number of DEGs between the two macaque species, some of which are associated with human neurogenic diseases (e.g., *ATP1B2*, *PDE11A*, and *NCR3LG1*). In addition, we find that the gene expression differences relevant to metabolisms between the two macaque species (e.g., *SLC13A5*, *PLAAT2*, and *FABP1* in livers). These findings highlight the significance of understanding the divergence in gene expression, particularly in disease-associated genes, when deciding between Mfa and Mmu as models for specific neurological disorders or metabolism disorders.

In addition to our comparative transcriptome analysis between the two macaque species, we also examine the comparative transcriptome analysis between macaques and humans. We find that more than ∼49.0% of one-to-one orthologous genes are expressed coincidently between humans and macaques. Tissue-specific genes are highly conserved between humans and macaques. Thus, these results underscore the substantial overlap in biological processes between the two species and support the merit of utility of macaques as biomedical models[34]. However, brain tissues exhibit the highest number of DEGs between humans and macaques and this discrepancy highlights potential differences in the evolutionary development of brains in primates[35]. These findings contribute to a more comprehensive understanding of the intricate interplay between gene expression, evolution, and the development of neurological traits in humans and macaques.

Building upon prior research showing the burst of segmental duplications in great apes[36, 37], we examine the relationship between structural variation and DEGs. Intriguingly, we find that 2,601 genes (∼19.41%) overlapped with lineage-specific structural variants are more prone to have gene expression differences. This observation suggests that SVs are one of major contributor to gene regulation, potentially driving the expression disparities observed[38]. One of particular interest, the case of *FAM220A*, underscores this correlation between copy number variation and gene expression between humans and macaques.

Our study provides a preliminary perspective on the comparative transcriptomes of two macaque species, as well as between macaques and humans. However, to obtain a more comprehensive understanding of macaque transcriptomes, it is imperative to expand sample size and encompass a broader range of tissues[39]. Moreover, while bulk RNA-sequencing has been widely used, it still grapples with the challenge of revealing the intricate cellular heterogeneity within tissues[40]. Additionally, the limitations of short-read RNA sequencing prevent us from capturing the full spectrum of isoform information within transcripts[41, 42]. In light of these challenges, the incorporation of long-read sequencing technology and single-cell RNA sequencing holds immense promise for unravelling the intricacies of macaque transcriptomes.

## Method

### Collection of animal tissues

A total of 8 female Mfa and 8 female Mmu were involved in this study. The use and care of animals complied with the guideline of Center for Excellent in Brain Science and Intelligence Technology. The animal tissues used in this study were collected from naturally deceased and euthanized monkeys for teaching purposes, in accordance with ethical guidelines. It’s worth noting that most of the tissues used in the study originated from the same individuals, with the exception of blood samples. The blood samples were collected from six healthy monkeys of similar age. To ensure the integrity of the samples, tissue collection was carried out within a maximum of five hours after death. Following collection, the samples were immediately snap-frozen in liquid nitrogen and stored in a liquid nitrogen environment until RNA extraction was performed.

### RNA-seq library preparation and sequencing

84 samples from fourteen different tissues were collected for Illumina next-generation sequencing. Total RNA was extracted with Trizol Reagent and qualified using NanoDrop. mRNA was purified from 3μg qualified total RNA. After fragmentation, first and second strand cDNA were synthesized. Adapters were ligated to cDNA fragments of 400-500 bp length purified by AMPure XP system. The final library was PCR enriched, quantified by Bioanalyzer 2100 system (Agilent), and sequenced on NovaSeq 6000 platform (Illumina).

### RNA-seq samples in humans

The normalized gene expression data (TPM) of all human RNA-seq samples were previously analyzed uniformly by the GTEx (v8) consortium and obtained from https://gtexportal.org/home/ datasets[43]. A total of 84 human samples corresponding to the 14 tissues of macaque samples were used. We obtained 15,000 one-to-one orthologous genes between human and rhesus macaque using the biomart tool in the Ensembl (v110). Considering the highly similar gene expression profiles across five brain tissues, including neocortical areas (prefrontal and visual cortices) and subcortical regions (striatum, hippocampus and thalamus), we consolidated them into one aggregated brain tissue group for subsequent integrated analysis.

### RNA-seq data processing

A total of 380 billion raw reads obtained from the eighty-four libraries we used for analysis. First, we used fastp (version 0.22.0) to control the quality of the raw reads and generate clean reads[44]. All the clean reads were then mapped to the reference genome of rheMac10(M. mulatta) using hisat2(version 2.2.0) with parameters ‘--dta --new-summary -p 10 -x’[13, 45]. Aligned reads were converted to BAM files and sorted based on genomic position using samtools (version 1.6.0)[46]. We also used samtools to remove the mitochondrial RNA contained aligned BAM files. Then the BAM files were counted for each annotated gene (n=33960) into a count matrix by R package Rsubread (version 2.12.0) with function featurecount[47]. We used R package edgeR with function DGEList and rpkm to normalize the count matrix[48]. Subsequently, TPM values were calculated from FPKM using the formula log(FPKM) -log(sum(FPKM)) + log(1e6), which accounts for gene length and library size.

### Sample clustering and quality control

In order to validate the reproducibility of individuals within each tissue type and to examine outliers with substantial within-group variations, we calculated the correlation of gene expression between samples. Ultimately, 84 samples with similar gene expression patterns within each group and minimal individual variations were selected for subsequent analyses, including 41 Mfa samples and 43 Mmu samples. Afterwards, we calculated the median gene expression in each tissue in Mfa and Mmu, respectively, to represent the gene expression value in specific tissues. Then, hierarchical clustering of median expression correlation matrix was performed using pheatmap (version 1.0.12)[49], to delineate the relationships between tissues of Mfa and that of Mmu based on median expression values. In addition, we performed dimensionality reduction on the gene expression of different samples using PCAtools (2.8.0) with parameter ‘mat = expression, metadata = sample_df, removeVar = 0.1’. We also used pheatmap package to delineate the relationships between tissues of human and two macaques based on median expression values, similarly as described above[50].

### Differential gene expression and tissue-specific gene analysis

To identify tissue-specific genes, raw count data of samples in each tissue were compared to those in other tissues using DESeq2 (version 1.36.0)[51]. We also used DESeq2 package to detect species-specific genes between Mfa and Mmu. DESeq2 calculated adjusted p-values using the Benjamini-Hochberg method to reduce false positive rate (FDR). Here, we utilized a log2 fold change (log2FC) greater than 1.5 and a statistical significance level of false discovery rate (p-adjust) less than 0.05 as the threshold criteria to identify both tissue-specific genes and species-specific genes. We performed hypergeometric tests to evaluate the significance of overlaps between tissue-specific genes from the two species using the phyper function, and adjusted the p-values using the Benjamini-Hochberg method. Gene ontology analysis for DEGs was performed using the ‘enrichGO’ function of the clusterProfiler with parameters ‘pAdjustMethod = ‘BH’, ont = ‘BP’, pvalueCutoff = 0.05, qvalueCutoff = 0.05,)’[52].

Differential gene expression analysis between humans and macaques across 10 tissues was conducted utilizing the limma (version 3.52.4) package[53], since the obtained human TPM matrix from the database was pre-normalized, not permitting further normalization through DESeq2. The TPM matrix comprising 15,000 human-macaque orthologous genes underwent analysis via the function model.matrix, lmFit, contrasts.fit, eBayes, and topTable functions in limma package. This analysis identified tissue-specific genes in both humans and macaques, alongside species-specific genes. For detecting tissue-specific genes, a threshold of log2(fold change) > 1.5 and false discovery rate (FDR) < 0.05 was utilized. In contrast, a more stringent threshold of log2(fold change) > 1.5 and FDR < 0.001 was utilized for identifying species-specific genes, to minimize intra-group variances and select for genes exhibiting higher significance. The rest of the analyses followed similar procedures aforementioned.

### Intersection of DEGs and Macaque-Specific SVs

We downloaded the macaque-specific structural variants with respect to the human genome (hg38) from public databases. We also generated summary data of DEGs in at least 2 tissues. The overlap between the gene set and structural variants (SVs) was examined with bedtools (version 2.30.0)[54]. Subsequently, a permutation test was conducted to examine the significance of this observation. Firstly, we generated 5,000 sets of randomly permuted structural variants (SVs) while preserving original SV lengths with the shuffle function in bedtools. Then, we used bedtools to intersect the genes overlapped with the macaque-specific SVs. Finally, p-values representing the statistical significance of our findings were derived by ranking the observed values and permuted values.

### Gene expression validation with qPCR experiment

Briefly, total RNA was extracted using Trizol Reagent following the manufacturer’s instructions and quantified by NanoDrop 1000 (Thermo Fisher). Gene expression levels were measured with Roche 480II Real-Time PCR System (Roche). Quantitative PCR was then carried out with specific primers for target genes, using *GAPDH* as an internal control. Finally, the relative expression levels were determined via the 2>(−ΔΔCt) method (Livak Method)[55]. The qRT-PCR primers are listed in the Supplementary Data (Table S10).

### Evolutionary history of FAM220A analysis

We downloaded the public copy number variation summary data from public database. Then, we used bedtools and custom scripts to select the genes which have copy number variation between humans and macaques, and have gene expression differences in at least two tissues. We also used minimiro to determine the syntenic relationship between apes and macaque in *FAM220A* locus. We extracted *FAM220A* region and used mafft to algin the sequences. Finally, we used BEAST2 to examine the time of duplicated events[56].

### Statistical analysis

Statistical analysis was performed using R software (v.4.2.3) (https://www.R-project.org/). Most of the figures in this paper were generated using ggplot2[57]. Adjusted p < 0.05 was considered significant for all the analysis.

## Data availability

The raw RNA-seq data reported in this study are available in NCBI with BioProject PRJNA1004471 and Sequence Read Archive accession numbers: SRR25608803-SRR25608886.

## Acknowledgements

We thank the Shanghai Songjiang Non-Human Primate Research Platform for providing all the macaque samples, and the veterinarians for collecting the samples. The computations in this paper were run on the cluster supported by Center for Data and Computing in Brain Science (CDCBS) at Institute of Neuroscience, Chinese Academy of Sciences; and on the Siyuan-1 and π 2.0 cluster supported by the Center for High Performance Computing at Shanghai Jiao Tong University.

## Funding

This work was supported by grants from the National Natural Science Foundation of China Grant (31825018, 82021001.), the National Key Research and Development Program of China (2022YFF0710901), and the National Science and Technology Innovation 2030 Major Program (2021ZD0200900) to Q. S. This work was supported by Shanghai Pujiang Program (22PJ1407300) and Shanghai Jiao Tong University 2030 Program (WH510363001-7) to Y.M.; and by National Natural Science Foundation of China (82001372) to X.Y.

### Contributions

Y.M., and Q.S. conceived the project. Y.X.M. performed the RNA-seq analysis. Y.L. and N.X. helped sample collection and RNA-sequencing. Z.Y., S.Z., X.Y. and X.W. performed the analysis of *FAM220A*. Y.X.M., X.Y., and N.X. performed the qRT-PCR experiments.

Y.X.M. and X.Y. contributed to the editing of the main image. Y.X.M., Q.S., and Y.M. drafted the manuscript. Y.X.M., Q.S., and Y.M. finalized the manuscript. All authors read, edited and approved the manuscript.

## Notes

### Competing Interest Statement

The authors have declared no competing interest.

